# Contributions of m6A RNA methylation to germline development in the planarian *Schmidtea mediterranea*

**DOI:** 10.1101/2025.06.02.657537

**Authors:** Junichi Tasaki, Labib Rouhana

## Abstract

*N*^6^-methyladenosine (m^6^A) is one of the most prevalent post-transcriptional modifications of eukaryotic RNA molecules. This post-transcriptional modification is essential in biological contexts ranging from metabolism to cellular differentiation and neuronal function. While the role of m^6^A RNA regulation in the soma of planarian flatworms has been previously studied, the presence and biological relevance of this regulatory pathway in the germline of these or other lophotrochozoans remains unknown. Here, we characterize m^6^A RNA regulation factors in the planarian *Schmidtea mediterranea* and analyze their function in development of the germline. Enriched expression of orthologs of m^6^A methyltransferase complex components, namely *Smed-METTL3*, *Smed-METTL14*, and *WTAP*, was detected in ovaries and testes of *S. mediterranea*. Perturbation of these factors by RNA-interference (RNAi) disrupted intermediate steps in oogenesis and spermatogenesis, but the presence of germline stem cells was not affected. Expression of m^6^A “reader” homologs was also detected in the planarian gonads. Sperm development defects were observed upon *Smed-YTHDF2-3* RNAi, and combined knockdown of *Smed-YTHDF2-3, Smed-YTHDF2-2*, or *Smed-YTHDF1-2* with that of writers led to increased frequency of germline development defects. These results indicate that regulation of RNA bym^6^A methylation supports male and female germline development in planarian flatworms.

## INTRODUCTION

*N*^6^-methyladenosine residues (m^6^A) are one of the most abundant modifications in eukaryotic transcripts, as these account for as much as 0.2-0.6% of all adenosines in RNA [1–3]. Deposition of m^6^A leads to alterations in RNA structure [4], and influences nuclear processing and export of mRNA [5, 6], as well mRNA stability [7] and translation [8]. The presence of this reversible regulatory mark is orchestrated by competition between a methyltransferase complex [9–11] and demethylase enzymes [6, 12] known as “writers” and “erasers”, respectively (reviewed by [13]). The writer complex contains the catalytic subunit Methyltransferase-Like 3 (METTL3), which cooperates with METTL14 for substrate recognition [7, 14] and with the regulatory subunit Wilms’ Tumor 1-Associated Protein (WTAP) [15, 16]. Demethylation of adenosine occurs passively, or through eraser demethylase proteins such as Fat-mass and Obesity-associated protein (FTO) and Alpha-ketoglutarate-dependent Dioxygenase Homologue 5 (ALKBH5) [6, 12]. When present, the m^6^A mark is recognized by “readers” such as YTHDF1, YTHDF2, and YTHDF3, which belong to YT521-B homology domain family of proteins. Readers recruit downstream regulators of mRNA to secure accurate regulation of targets and progression of biological processes [7, 17–20].

Proper regulation of m^6^A dynamics is essential for development and viability of plants and animals [7, 21–25]. Inhibition of m6A-associated machinery has been shown to affect several developmental processes, including stem cell differentiation in vertebrate embryos [7, 23, 26], sex determination in flies [24, 27], and development of the germline in *Drosophila* and in mammals [6]. In flies, the catalytic subunit of the m^6^A methylation complex regulates oogenesis by modulating Notch signaling [28]. In mammalian testes, ablation of *Mettl3* severely inhibits spermatogonia differentiation and initiation of meiosis [29], whereas increased m^6^A abundance leads to apoptosis of meiotic metaphase stage spermatocytes [6]. These and other studies support the hypothesis that regulation of RNA by m^6^A based processes plays a role in cellular differentiation. Whether m^6^A machinery regulates conserved cellular differentiation processes or through regulation of homologous targets remains to be determined.

Planarians are free-living freshwater flatworms that are members of the phylum Platyhelminthes (superphylum Lophotrochozoa; reviewed by [30, 31]). Planarians have bilateral symmetry, derivatives of all three germ layers, and distinct digestive, nervous, nephridial, and reproductive systems [32–37]. Planarians can reproduce asexually by transverse fission and regeneration, or sexually as cross-fertilizing hermaphrodites, although some species only exhibit one reproductive mode [38, 39]. Classically, planarian flatworms have been studied mainly for their capacity of full-body regeneration [40, 41]. However, there is a growing interest in studying germline development in planarians, as formation of the entire reproductive system can be observed post-embryonically and is derived from adult pluripotent stem cells known as neoblasts [33, 42].

Conserved components of post-transcriptional regulation machineries present in germ granules, such as Vasa, Tudor, and PIWI homologs, have been shown to regulate maintenance and differentiation of planarian neoblasts [43–52]. A subset of conserved RNA-binding proteins, which include Nanos [53, 54], Boule [55], CPEB [56], Bic-C [57], and one particular PIWI [58], have been shown to be specifically expressed and required during development of the planarian germline. Hence, the transition from neoblasts to germline stem cells to gametes seems to involve a progression of subtle changes in post-transcriptional regulation programs. The involvement of RNA regulation via m^6^A modification in the developmental progression from neoblasts to gametes remains to be determined.

In this study, we assess the presence and function of m^6^A writers, readers, and erasers in *Schmidtea mediterranea* planarian hermaphrodites. Biological functions of m^6^A were analyzed during planarian germline development and regeneration. Expression of methyltransferase genes, *Smed-METTL3* and *Smed-METTL14*, was detected in the nervous system, ovaries, testes, and germline stem cells in sexual animals. Both *Smed-METTL3* and *Smed-METTL14* were also expressed in the nervous system in asexual animals. Perturbation of the m6A pathway using RNAi did not affect maintenance of germline or pluripotent somatic stem cells. However, RNAi targeting the writers, *Smed-METTL3* and *Smed-METTL14*, resulted in oocyte and sperm development defects. In addition, homologs of YTHDF m^6^A-binding proteins were expressed in the same anatomical regions as the methyltransferases in *S. mediterranea*. RNAi targeting expression of specific readers resulted in germline development defects or exacerbated defects observed upon knockdown of writer genes. Collectively, these results extend previous studies of m^6^A RNA regulation in asexual planarians [59–62] by providing evidence of its contribution in male and female germ cell development in *S. mediterranea* hermaphrodites.

## RESULTS

### m^6^A RNA methylation writer and reader gene homologs are expressed in the brain and gonads of *Schmidtea mediterranea*

Human sequences of METTL3 and METTL14 methyltransferases, WTAP, demethylases ALKBH5 and FTO, and m^6^A-binding proteins (YTHDF1, YTHDF2 and YTHDF3) were used as input to identify homologs in *S. mediterranea*. A TBLASTN search against reference transcriptome sequences in the PlanMine database [63, 64], revealed matches for METTL3, METTL14, WTAP, and m^6^A-binding proteins YTHDF1 and YTHDF2 in *S. mediterranea* (Table 1). Potential orthologs of demethylases, ALHBH5 and FTO, were not found using this approach. Identified m^6^A methylation pathway homologs matched individual genes in the *S. mediterranea* genome (SmedGD; [65]) and were named *Smed-METTL3*, *Smed-METTL14*, *Smed-WTAP*, *Smed-YTHDF1-1*, *Smed-YTHDF1-2*, *Smed-YTHDF2-1*, *Smed-YTHDF2-2*, and *Smed-YTHDF2-3* to follow the nomenclature used by Dagan et al. [59] (Table1). As indicated in the list of identified genes and assigned names (Table 1), we found planarian genes coding for proteins with closest sequence identity to human YTHDF1 and YTHDF2. However, potential orthologs of *YTHDF3* were not identified. These results indicate that m^6^A writing and reading machineries are present in *S. mediterranea*, whereas erasers appear to have been lost.

**Table 1.** m^6^A genes characterized in sexual *S. mediterranea*. Gene names of *S. mediterranea* homologs of m^6^A pathway components following nomenclature used by Dagan et al. (first column), along with reference identifiers in the dd_Smed_v6 (second column) and SMES (third and fourth columns) transcript designations. Top match human reference protein in BLASTX analyses (fifth column), along with match coverage, E-value, and percent identity (sixth, seventh, and eight columns) are also shown.

| *S. mediterranea* Gene* | dd_Smed_v6 ID | SMES Transcript ID | Redundant Transcript IDs | BLASTX *H. sapiens* REF_SEQ_SELECT Top Hit | Coverage | E value | Identity |
| --- | --- | --- | --- | --- | --- | --- | --- |
| *METTL3* | dd_Smed_v6_6450_0_2 | SMEST034548001.1 |  | METTL3 - methyltransferase 3, N6-adenosine-methyltransferase complex catalytic subunit | 68% | 1.00E-180 | 56.28% |
| *METTL14* | dd_Smed_v6_6641_0_1 | SMEST073528001.1 |  | METTL14 - methyltransferase 14, N6-adenosine-methyltransferase non-catalytic subunit | 51% | 3.00E-143 | 69.21% |
| *WTAP* | dd_Smed_v6_2322_0_4 | SMEST043489001.1 | SMEST043668001.1 | WTAP - WT1 associated protein | 23% | 2.00E-41 | 44.12% |
| *ythdf1-1* | dd_Smed_v6_7162_0_1 | SMEST039495001.1 |  | YTHDF1 - YTH N6-methyladenosine RNA binding protein F1 | 23% | 3.00E-58 | 54.55% |
| *ythdf1-2* | dd_Smed_v6_8450_0_1 | SMEST003195002.1 | SMEST003195001.1 | YTHDF1 - YTH N6-methyladenosine RNA binding protein F1 | 26% | 7.00E-46 | 47.59% |
| *ythdf2-1* | dd_Smed_v6_5316_0_1 | SMEST075874001.1 |  | YTHDF2 - YTH N6-methyladenosine RNA binding protein F2 | 36% | 3.00E-56 | 52.57% |
| *ythdf2-2* | dd_Smed_v6_5578_0_1 | SMEST001953001.1 |  | YTHDF2 - YTH N6-methyladenosine RNA binding protein F2 | 24% | 2.00E-56 | 52.38% |
| *ythdf2-3* | dd_Smed_v6_7891_0_1 | SMEST008619001.1 |  | YTHDF2 - YTH N6-methyladenosine RNA binding protein F2 | 26% | 8.00E-80 | 68.36% |
| *ythdc-1\*\** | dd_Smed_v6_3491_0_1 | SMEST018260001.1 |  | YTHDC1 - YTH N6-methyladenosine RNA binding protein C1 | 57% | 3.00E-63 | 45.56% |

\*Names based on Yael Dagan et al., 2022
\*\*Not analyzed in this study.

We predicted that genes responsible for m^6^A-mediated regulation of mRNA would share similar distribution patterns of expression. Thus, we assessed the distribution of expression of the eight potential m^6^A pathway components by whole-mount *in situ* hybridization (WMISH) in asexual and sexual strains of the planarian *S. mediterranea*. Expression of *Smed-METTL3*, *Smed-METTL14*, *Smed-WTAP*, *Smed-YTHDF1-1*, and *Smed-YTHDF2-3* was detected in the brain of asexual planarians, which can be observed as a horseshoe structure in the region surrounding the eyes (Fig. 1, A-E). Expression of *Smed-YTHDF2-2* and *Smed-YTHDF1-2* were detected in the planarian gut (Fig. 1, F and G), whereas *Smed-YTHDF2-1* expression was not conclusively detected in any specific structure (Fig. 1H). Some expression signal was detected in the gut for *Smed-METTL3*, *Smed-METTL14*, and *Smed-WTAP* samples (Fig. 1, A-C). Given that background signal in the gut is often observed in planarian samples analyzed by colorimetric WMISH, we reassessed the distribution of expression *Smed-METTL3* and *Smed-METTL14* by fluorescence *in situ* hybridization (FISH). Expression of *Smed-METTL3* and *Smed-METTL14* in the brain was validated using FISH, which also revealed some signal in the ventral nerve chords, but not in the gut (Fig. 1, A’ and B’). Given these results, we concluded that the m^6^A writer complex and at least two readers (*YTHDF1-1* and *YTHDF2-3*) are preferentially expressed in the planarian brain of asexual planarians.

**Figure 1.**
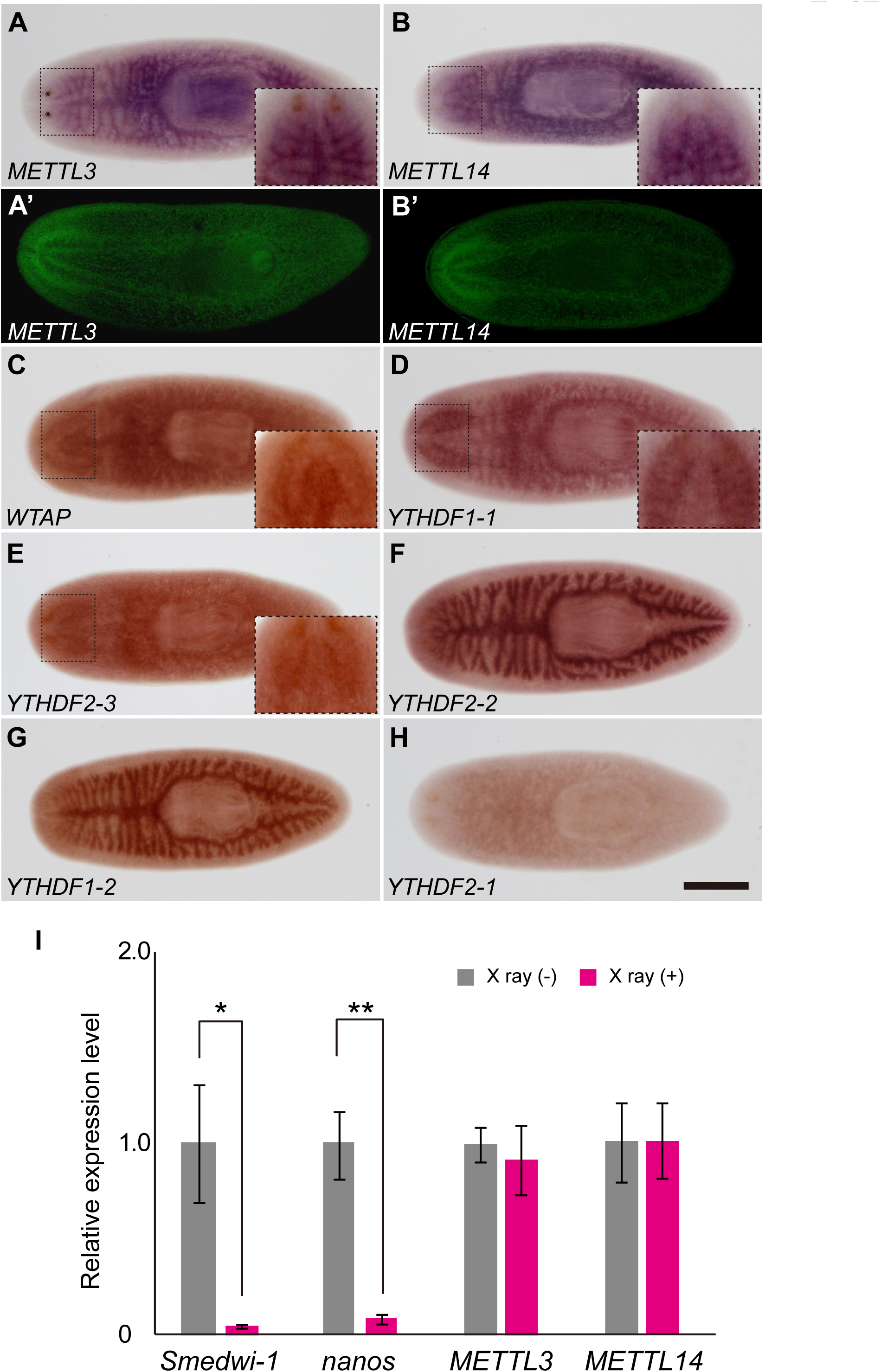
Expression pattern of m^6^A pathway genes in the asexual strain. **(A-H)** Colorimetric whole-mount *in situ* hybridization (A-H) and FISH (A’-B’) reveal enriched expression of *Smed-METTL3* (A and A’) and *Smed-METTL14* in the *S. mediterranea* nervous system. Insets show close-up of brain region. Enriched expression in the brain was also observed for *Smed-WTAP*, *Smed-YTHDF1-1*, and *Smed-YTHDF2-3* (C-E). Enriched expression of *Smed-YTHDF2-2* and *Smed-YTHDF1-2* (F-G) was detected in intestinal cells. No conclusive signal of *Smed-YTHDF2-1* expression was detected (H). (I) RT-qPCR analysis of gene expression in intact (gray) and x-ray irradiated (magenta) planarians shows ablation of expression of the neoblast marker *Smedwi-1* and the germline stem cell marker *nanos* 7 days post-irradiation, whereas relative expression levels of *Smed-METTL3* and *Smed-METTL14* were not significantly changed by x-ray irradiation. Relative expression to control samples is shown after normalizing gene-specific RT-qPCR level to that of *ß-tubulin* in each sample. Scale bar: 500 μm.

Expression of m^6^A writer and reader genes in neoblasts was not evident from *in situ* hybridization analyses. To go a step further in assessing the possibility that this regulatory mechanism contributes to the biology of planarian stem cells, neoblasts were depleted from asexual planarians by X-ray irradiation and expression of *Smed-METTL3* and *Smed-METTL14* was compared between intact control and irradiated asexual planarians by reverse transcription quantitative PCR (RT-qPCR). The expression *Smedwi-1* and *nanos* were measured as indicator of neoblast [47] and germline stem cell [53] depletion efficiency, respectively, seven days after irradiation. As expected, the abundance of *Smedwi-1* and *nanos* mRNAs dropped by 90% or more in planarians subjected to X-ray irradiation (Fig. 1I). On the other hand, the abundance of *Smed-METTL3* and *Smed-METTL14* transcripts was not significantly different upon neoblast depletion (Fig. 1I), providing no evidence of *Smed-METTL3* or *Smed-METTL14* expression in planarian adult somatic stem cells. We consulted single-cell RNA-seq (scRNA-seq) data of Plass et al. [66] and Zheng et al. [67], which revealed some expression of *Smed-METTL3* and *Smed-METTL14* in neoblasts, and peak levels of expression during neuronal differentiation. These results indicate that *S. mediterranea* homologs of m^6^A writer complex genes are preferentially expressed in the nervous system in asexual animals.

To assess the possibility that m^6^A regulation plays a role in planarian germline development, we performed WMISH analyses in adults of the sexual strain of *S. mediterranea* (Fig. 2, A-I). In addition to corroborative detection of expression of *Smed-METTL3*, *Smed-METTL14*, *Smed-WTAP*, and *Smed-YTHDF1-1* in the brain (Fig. 2, A-D), expression of some of these genes was also detected in gonads. More specifically, expression of *Smed-METTL3*, *Smed-METTL14*, and *Smed-WTAP* was detected in testes and ovaries (Fig. 2, A-C and A’-C’). Furthermore, faint detection of *Smed-YTHDF2-3* (Fig. 2E) and *Smed-YTHDF2-2* (Fig. 2F) expression was detected in testes. Expression of *Smed-YTHDF1-2* was detected in the region of the ovaries (Fig. 2G). Expression of S*med-YTHDF2-1* was not detected in any specific organ of sexual planarians (Fig. 2H), as was the case during analysis of asexual samples (Fig. 1H). Altogether, these expression analyses revealed three fundamental subunits of the m^6^A methylation complex and at least one reader are present in the gonads and brain of *S. mediterranea*, suggesting the possibility that planarian germ cells rely on m^6^A regulatory pathways for their development and/or function.

**Figure 2.**
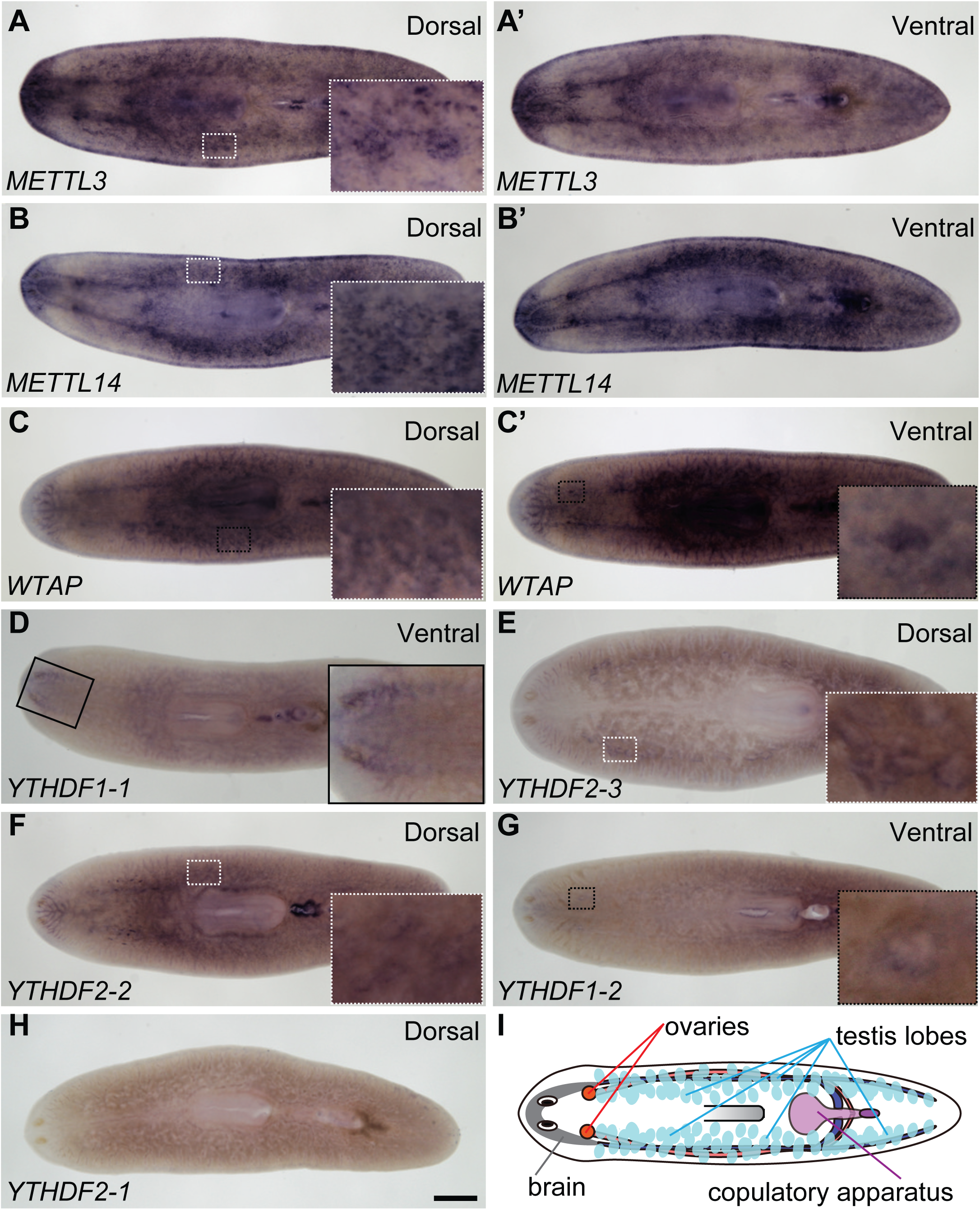
Expression of m^6^A pathway genes in gonalds of *S. mediterranea* hermaphrodites. **(A-C)** Expression of *METTL3* (A-A’), *METTL14* (B-B’) and *WTAP* (C-C’) is detected in testes (A-C, insets), ovaries (A’-C’, inset in C’) and brain (A’-C’) of sexual planarians using colorimetric whole-mount *in situ* hybridization. **(D-H)** Analyses m^6^A reader genes reveal enriched expression of *YTHDF1-1* (D; inset) in the brain, *YTHDF2-3* (E; inset) and *YTHFD2-2* (F; inset) in testes, and *YTHDF1-2* (G; inset) expression in ovaries. *YTHDF2-1* (H, I) expression was not conclusively detected. **(I)** Illustration depicting planarian anatomy. Scale bar: 500 μm.

### *Smed-METTL3* and *Smed-METTL14* are co-expressed in ovarian germline stem cells and oocytes of *S. mediterranea*

To determine whether m^6^A deposition could occur in germ cells, we performed detailed expression analysis of methyltransferase complex subunits by FISH and confocal microscopy in ovaries (Fig. 3, A-C). Expression of *Smed-METTL3* was observed in cells co-expressing the germline stem cell marker *nanos* in the periphery of the ovary, and prevailing in mature oocytes (Fig. 3, A-A”). Expression of *Smed-METTL14* was also observed in germline stem cell and oocytes of planarian ovaries (Fig. 3, B-B”). Furthermore, co-detection of *Smed-METTL3* and *Smed-METTL14* transcripts revealed overlapping expression throughout the ovary and abundantly in fully grown oocytes (Fig. 3, C-C”), providing support for the notion that m^6^A methyltransferase activity is present in the germline an involves the formation of METTL3/METTL14 heterodimers in planarians as in mammals [11].

**Figure 3.**
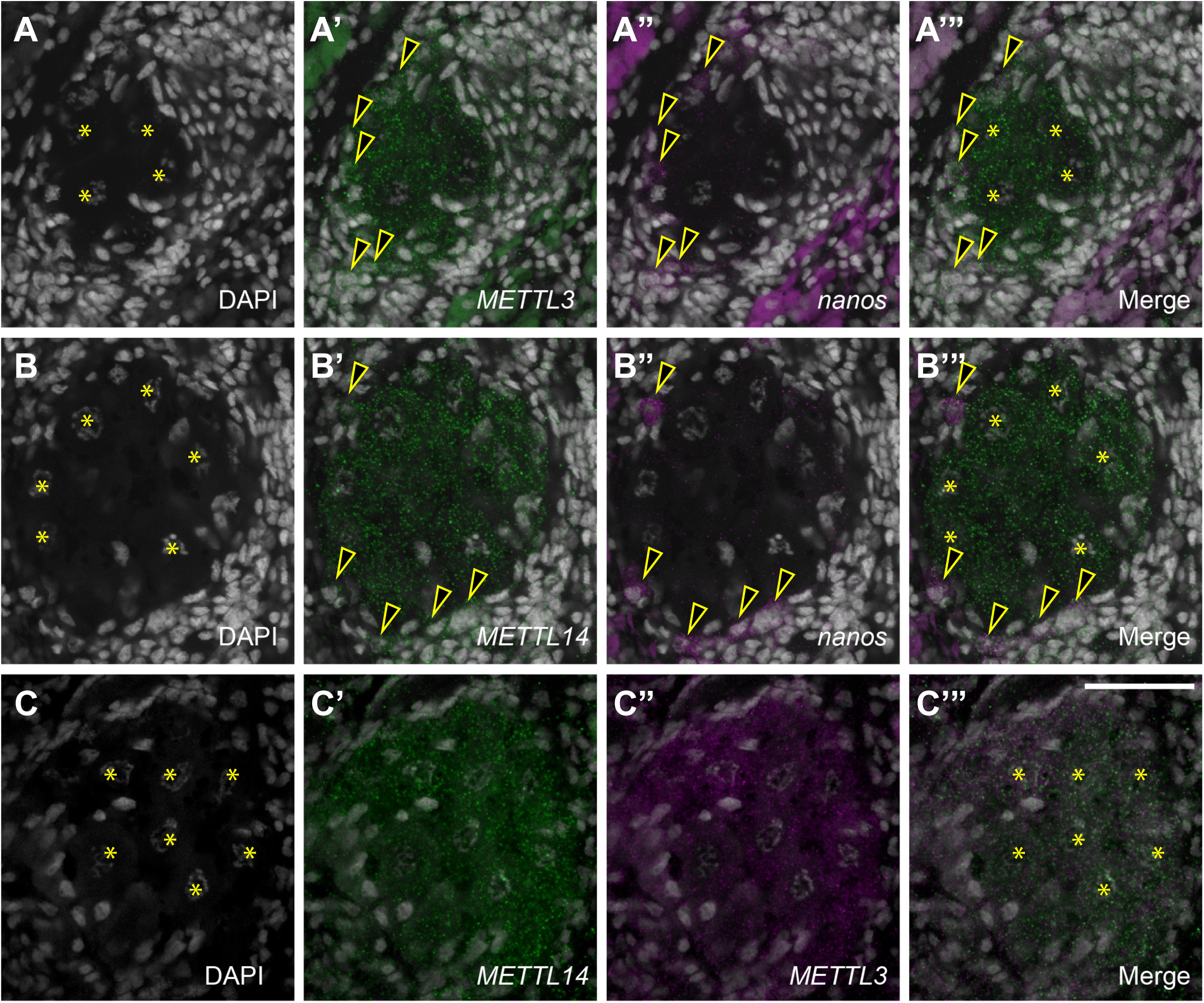
*Smed-METTL3* and *Smed-METTL14* are co-expressed in germline stem cells and oocytes of planarian ovaries. **(A-C)** Detection of double FISH signals in single plane confocal images reveal overlapping presence of *Smed-METTL3* (A’) and *nanos* (A”) mRNA in ovarian germline stem cells (arrowheads), while *Smed-METLL3* mRNA is also present in oocytes (asterisks in (A)). Double FISH analysis of *Smed-METTL14* and *nanos* expression (B-B”) as in (A-A”) reveals the presence of *Smed-METTL14* in ovarian germline stem cells and oocytes, while double FISH analysis of *Smed-METTL3* and *Smed-METTL14* (C-C”) confirms overlapping expression throughout the ovary. Cell nuclei are stained with DAPI (gray) in all panels. Scale bar: 50 μm.

### m^6^A RNA regulation contributes to normal development of male and female gonads in *S. mediterranea*

RNA interference (RNAi) was performed to assess the function of m^6^A methylation in sexual planarians. To do this, *S. mediterranea* hermaphrodites were subjected to a diet of calf liver containing double-stranded RNA (dsRNA) targeting either *Smed-METTL3* or *Smed-METTL14*. Ingestion of dsRNA induces systemic RNAi in planarians and is sustained by weekly feedings to allow cellular turnover from neoblasts under knockdown conditions [68]. A group of planarians were fed dsRNA corresponding to firefly *luciferase* sequence, which is known not to interfere with normal physiology or germline development in *S. mediterranea* [69, 70]. After six weeks of RNAi, the anatomy of gonads was analyzed by DAPI staining and confocal microscopy as per Wang et al. [57]. All of the control planarian samples displayed ovaries with fully developed oocytes (Fig. 4A; asterisks) and testis lobes with plentiful spermatozoa (Fig. 4E; arrows) indicating normal development of reproductive structures. In contrast, subsets of planarians from the groups subjected to *Smed-METTL3* RNAi (*METTL3(RNAi)*; Fig. 4, B and F) or *Smed-METTL14* RNAi (*METTL14(RNAi)*; Fig. 4, C and G) displayed underdeveloped female and male gonads that lacked spermatozoa and oocytes, respectively. The frequency of underdeveloped gonads and incomplete germline development phenotypes increased by simultaneous knockdown of *Smed-METTL3* and *Smed-METTL14* (*METTL3;METTL14(RNAi)*), resulting in 85% of the ovaries lacking oocytes (Fig. 4D) and 71% of samples with azoospermia (Fig. 4H). These results show that m^6^A methyltransferases are required for normal development of male and female gonads in *S. mediterranea*.

**Figure 4.**
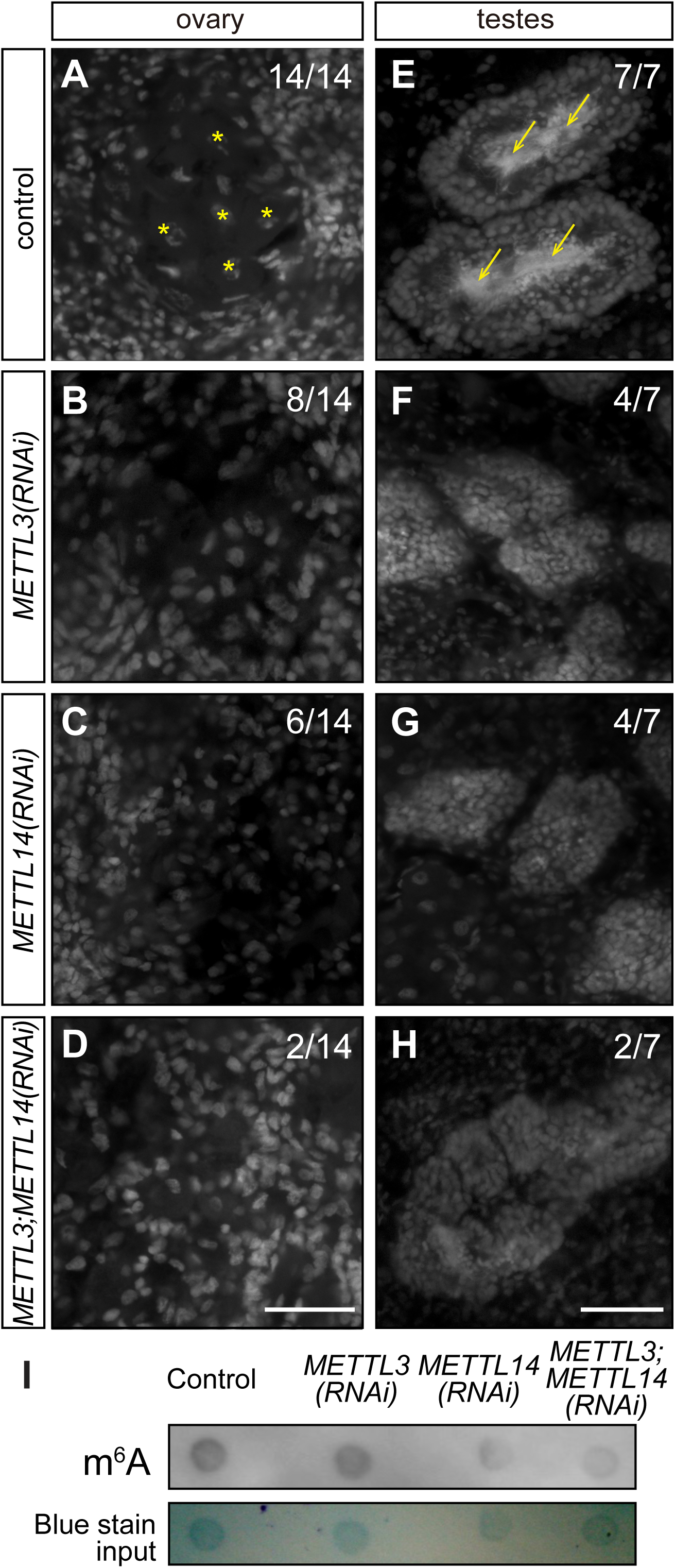
*Smed-METTL3* and *Smed-METTL14* function is required for normal development of ovaries and testes. **(A-D)** Confocal images of DAPI signal from ovaries of planarians show oocytes in ovaries of control knockdowns (asterisks in (A)), which are absent in ovaries of planarians subjected to RNAi against *Smed-METTL3* (B), *Smed-METTL14* (C), or *Smed-METTL3*;*Smed-METTL14* (D). **(E-H)** Sperm that populates the innermost region of testis lobes in control planarians (arrows; E) are not present in planarian knockdowns for *Smed-METTL3* (F) and for *Smed-METTL14* (G), and more frequently upon simultaneous knockdown of both genes (H). Parentheses show fraction of animals that displayed normal development of gonads. **(I)** Dot blot analysis showing decrease of m^6^A abundance detection (top) relative to total RNA (bottom) from *METTL3(RNAi)*, *METTL14(RNAi)*, and *METTL3;METTL14(RNAi)* animals as compared to controls. Scale bars: 50 μm

Given the gonad development phenotypes observed upon *Smed-METTL3*, *Smed-METTL14*, and *METTL3;METTL14* RNAi, we hypothesized that regulation via m^6^A deposition had been disrupted upon knockdown of components of the writer complex. To test this, m^6^A abundance was assessed by m^6^A antibody blot on total RNA extracts from knockdown planarians. The m^6^A signal levels were noticeably decreased in RNA from *METTL3(RNAi)* animals and *METTL14(RNAi)* animals, and most severely in RNA from *METTL3;METTL14(RNAi)* animals (Fig. 4I). These results show that m^6^A RNA abundance is dependent on normal expression of *Smed-METTL3* and *Smed-METTL14*, which indicates that regulation of gene expression via m^6^A RNA methylation is required for normal development of gonads in *S. mediterranea*.

### m^6^A RNA writers and readers support normal progression of male and female germline development in *S. mediterranea*

To gain insight into events that may be most sensitive to m^6^A RNA regulation during planarian germline development, we performed RNAi analyses in asexual and sexual planarians followed by FISH using stage-specific germ cell markers. In asexual planarians, neoblasts give rise to clusters of presumptive germline stem cells marked by expression of *nanos*, but these do not progress into later stages of gametogenesis [53]. Knockdown of methyltransferases, *Smed-WTAP*, or readers did not change the presence or distribution of presumptive germline stem cell clusters in asexual planarians (Supplementary Fig. S1). Likewise, there were no noticeable deficits on regeneration (Supplementary Fig. S2) or fission (Supplementary Fig. S3) observed in asexual planarians with disrupted expression of m^6^A methylation writers or readers during these analyses. These results suggest that, while m^6^A writers and readers may be expressed in asexual planarians, differentiation of neoblasts into germline stem cells are not sensitive to disruption of m^6^A regulation.

To further analyze the germline development defects caused by disrupting expression of m^6^A regulators, we performed combinatorial RNAi of readers and writers in sexual planarians followed by analyses by FISH. We hypothesized that knocking down readers expressed in male and female gonads would exacerbate the oogenesis and spermatogenesis defects observed upon knockdown of *METTL* genes. We also hypothesized that knockdown of the writer complex accessory protein WTAP would cause germline development defects similar to those observed upon disruption of expression of methyltransferases. As in our initial experiments (Fig. 4), all the samples from the control group (i.e. planarians treated with *luciferase* dsRNA) possessed readily detectable oocytes in their ovaries (Fig. 5A) and sperm in the innermost region of testis lobes (Fig. 6A). The presence of oocytes in these experiments was validated not by their morphology alone, but also by detection of the oocyte marker *Smed-CPEB1* [56] (Fig. 5A’). Similarly, the presence of germline stem cells and/or cells in early stages of gametogenesis in the gonads of control planarians (*e.g.*, oogonia and spermatogonia) were validated by detection of the *gH4* marker [53, 71] (Fig. 5A’ and 6A’). In contrast, simultaneous knockdown of *Smed-METTL3* and *Smed-METTL14* disrupted oogenesis (Fig. 5B) and spermatogenesis (Fig. 6B) in a subset of samples (Fig. 5G and Fig. 6I). However, cells expressing the *gH4* marker were detected in all the ovaries (Fig. 5B’) and testes (Fig. 6B’) of *Smed-METTL3*;*METTL14(RNAi)* samples that lacked gametes, indicating the presence of germ cells in early stages gametogenesis in these gonads. Loss of gametes was also observed in ovaries and testes of planarians subjected to disruption of *Smed-WTAP* expression alone (Fig. 5C-C’ and 6C-C’) or in combination with *Smed-METTL3* and *Smed-METTL14* (Fig. 5D-D’ and 6D-D’). Knockdown of *Smed-YTHDF1-2* expression, which was detected in ovaries (Fig. 2G), phenocopied oogenesis defects in planarians with disrupted expression of writer complex subunits (Fig. 5D-D’). The penetrance of this phenotype increased from 25% to 83% when *Smed-METTL3* and *Smed-METTL14* RNAi was combined with *Smed-YTHDF1-2* RNAi (Fig. 5F-F’ and 5G). Nevertheless, germ cells expressing *gH4* were observed in the gonads of all the samples (Fig. 5, A’-F’; Fig. 6, A’-H’). These results indicate that regulation of RNA via m^6^A deposition is required for progression of oogenesis and spermatogenesis in *S. mediterranea* and suggest that the reader homolog *Smed-YTHDF1-2* plays a pivotal role in mediating regulation of m^6^A RNA during oogenesis.

**Figure 5.**
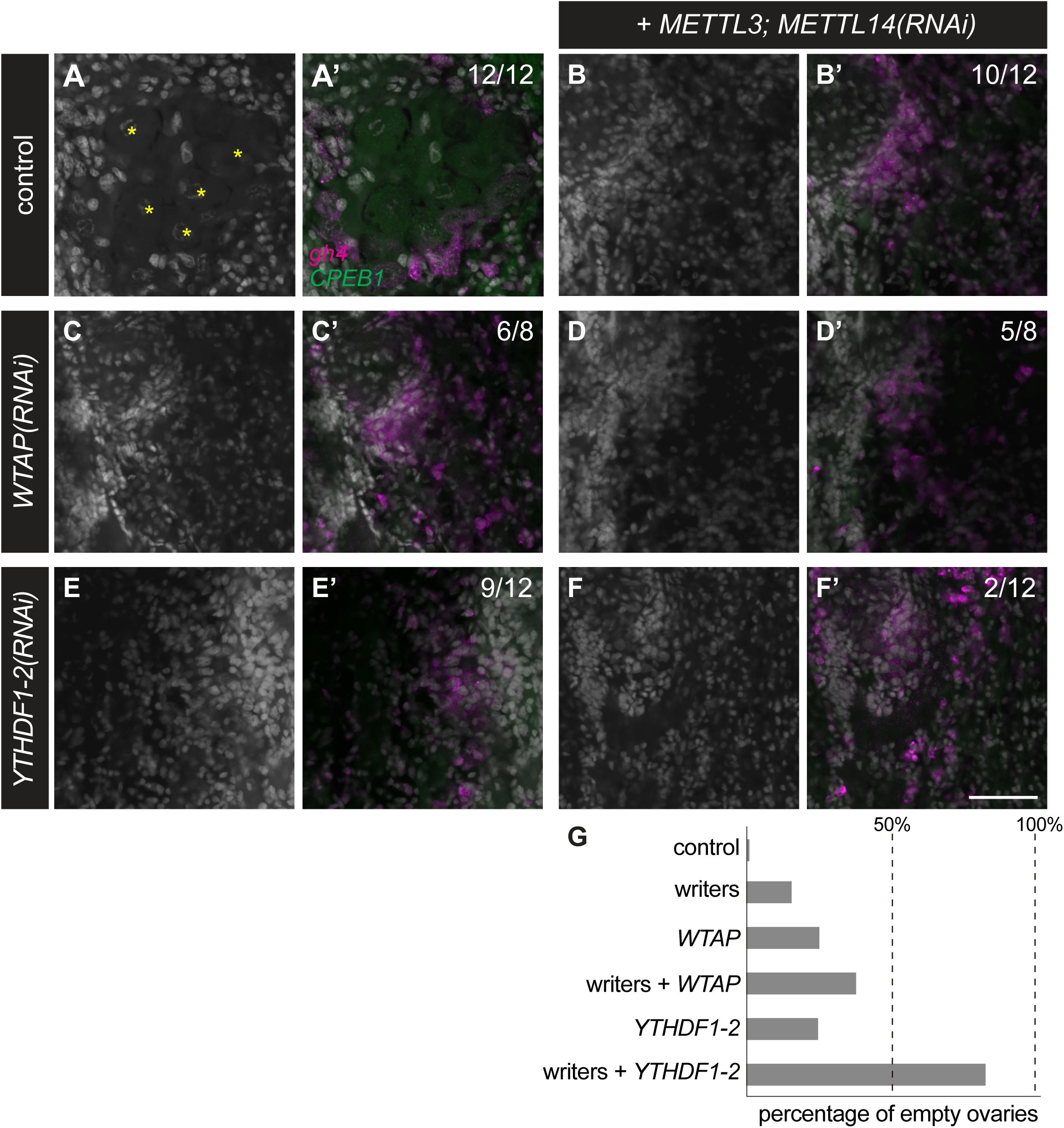
*Smed-YTHDF1-2* is required for normal oocyte development. **(A-F)** Confocal images of ovary region in control planarians (A-A’), and planarians subjected to knockdown of the writer genes (B), as well as *Smed-WTAP* or *Smed-YTHDF1-2* alone (C and E, respectively), or in combination with *Smed-METTL3*;*Smed-METTL14(RNAi)* (D and F). Cell nuclei stained by DAPI (A-F and A’-F’), as well as FISH detection of oocytes using the marker *CPEB1* (green) and of germ cells at earlier stages of oogenesis using *gH4* (magenta) (A’-F’) are shown. Knockdown of individual genes resulted in absence of oocytes from ovaries (C and E) which were observed in all control samples (marked by asterisks in (A)). The frequency of oocyte detection increased when knockdown of *Smed-YTHDF1-2* was combined with knockdown of writer genes (D). **(G)** Quantification of oogenic loss phenotype from experiments in (A-F). Scale bar: 50 μm.

**Figure 6.**
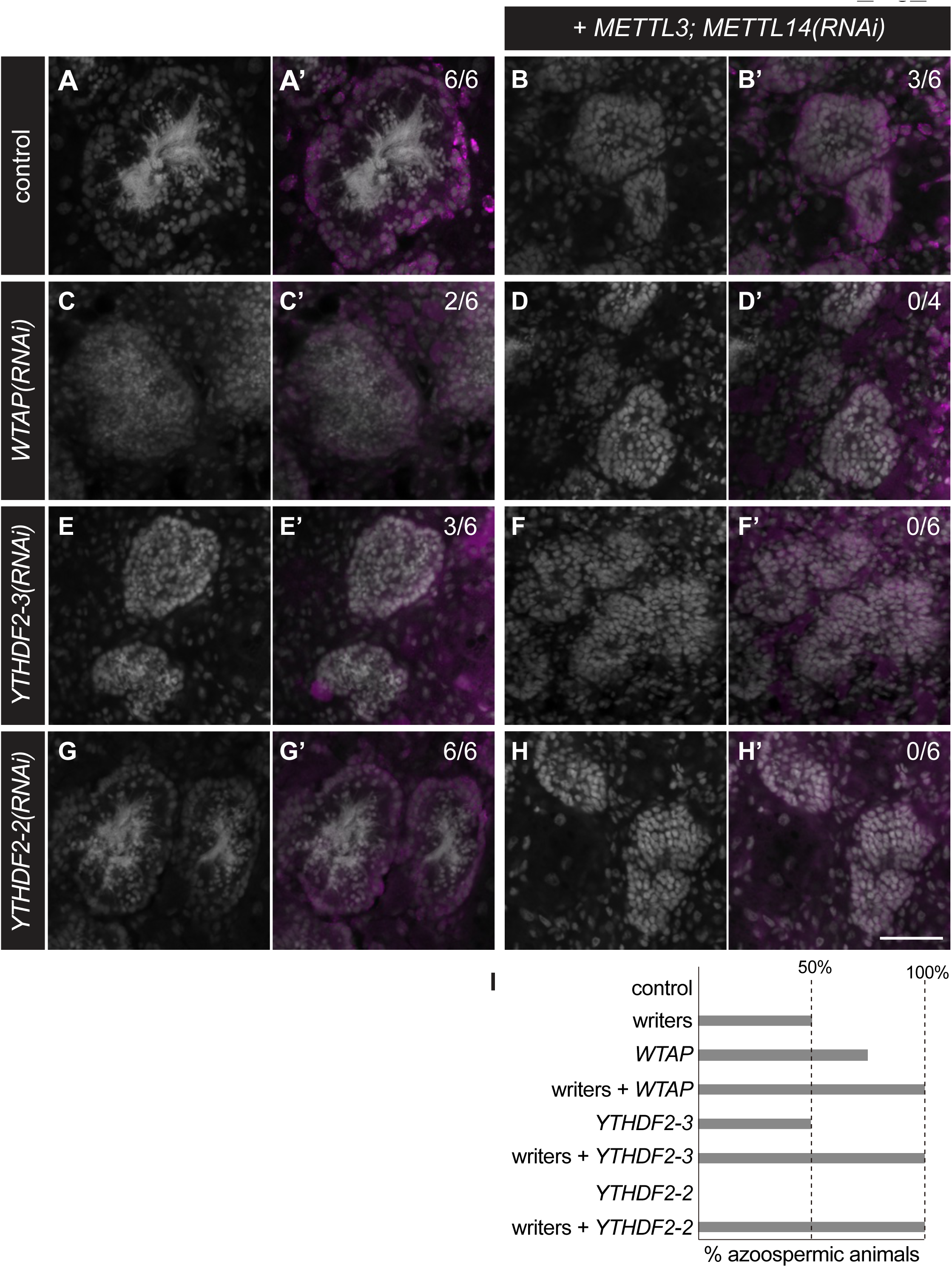
C *Smed-YTHDF2-3* and *Smed-YTHDF2-2* support sperm development. **(A-H)** Development of spermatozoa with elongated nuclei is observed by DAPI staining (white) in lumen of testis lobes from control (A) and *Smed-YTHDF2-2(RNAi)* (G) planarians but disrupted in a fraction of samples upon knockdown of writer genes (*Smed-METTL3;Smed-METTL14*) (B), *Smed-WTAP* (C) and the reader gene *Smed-YTHDF2-3* (E). The frequency of sperm loss increased to 100% of the samples upon simultaneous knockdown of *Smed-METTL3* and *Smed-METTL14* with *WTAP* (D), *Smed-YTHDF2-3* (F), and *Smed-YTHDF2-2* (H). Analysis by *gH4* FISH (magenta in A’-H’), which is a marker of germline stem cells and spermatogonia inside testis lobes as well as neoblasts in the parenchymal space (A’-H’) revealed that the presence of *gH4*-positive cells in testis lobes was constant in all samples. Fraction of samples with normal sperm development is shown in parenthesis for each treatment. (I) Quantification of loss of sperm phenotypes ins experiments represented by (A-H). Scale bar: 50 μm.

Expression of two reader homologs, *Smed-YTHDF2-3* and *Smed-YTHDF2-2*, was detected in testis lobes of sexual planarians by whole-mount *in situ* hybridization (Fig. 2E and 2F). We tested whether the function of these readers is required for spermatogenesis by disrupting their expression individually and in combination *Smed-METTL3*;*Smed-METTL14* RNAi (Fig. 6E-I). Disruption of *Smed-YTHDF2-3* expression alone caused spermatogenesis defects similar to those of *Smed-METTL3*;*Smed-METTL14(RNAi)* in half of the samples in the analysis (Fig. 6E-E’). The penetrance of this phenotype increased to 100% when *Smed-YTHDF2-3* knockdown was combined with *Smed-METTL3*;*Smed-METTL14* RNAi (Fig. 6F-F’ and 6I). In contrast, *Smed-YTHDF2-2* RNAi alone did not result in spermatogenesis defects (Figure 6G-G’ and 6I), but did increase the penetrance of the azoospermia to 100% of samples when combined with *Smed-METTL3*;*Smed-METTL14* RNAi (Fig. 6H-H’ and 6I). These data indicate that members of the writer complex, as well as Smed-YTHDF2-3 and YTHDF2-2, support sperm development in *S. mediterranea*. The detection of *gH4*(+) cells in the testes and ovaries of every animal displaying spermatogenesis and oogenesis defects in these experiments (Fig. 5B’-F’ and Fig. 6B’-6H’) corroborates with the continuity of germline stem cell clusters observed upon disruption of m^6^A writer and reader genes in asexual planarians (Supplementary Fig. S1). We hypothesize that regulation of mRNA by m^6^A is not required for development or maintenance of germline stem cells in planarian flatworms, but it is crucial for completion of spermatogenesis and oogenesis.

## DISCUSSION

This study characterized expression and function of the m^6^A methylation pathway in *S. mediterranea* sexual hermaphrodites. The data presented revealed expression of m^6^A writer complex components and readers in gonads of *S. mediterranea*, as well as their functional requirement for development of sperm and ova. These findings corroborate with studies in *Drosophila*, zebrafish, and mice that identified the requirement of m^6^A RNA methyltransferase machinery for completion of gametogenesis [28, 29, 72, 73]. The apparent conservation of m^6^A regulation during gametogenesis in planarians, flies, and mice supports the hypothesis that regulation of RNA through this post-transcriptional modification is ancestral to Bilateria, while evidence from studies of reproductive organs in *Arabidopsis* [21, 74] and meiotic cells in yeast [75] indicate that the function of m^6^A regulation during gamete development may be shared across eukaryotes.

Previous studies characterized components of the m^6^A RNA methylation machinery in asexual planarians [59–62]. Regeneration and growth defects were observed upon RNAi-mediated knockdown of components of the writer complex (*WTAP*, *KIAA1429* and *METTL14*) and one reader (*YTHDC*-*1*) in some of these studies [59, 60]. Unlike previous studies, we failed to detect defects in the somatic homeostasis or regeneration upon knockdown of m^6^A writer machinery, which may be due to differences in RNAi protocols (e.g. the use of higher amounts of dsRNA by Dagan et al. [59]). One collective observation amongst the current and previous studies is that neoblasts are viable in the absence of normal m^6^A RNA deposition. However, differentiation processes (whether of neoblasts into somatic cell types or during progression of spermatogenesis and oogenesis) require proper expression of m^6^A writer and reader machinery. The post-transcriptional control of cell cycle regulators by m^6^A, as observed in asexual planarians [60], may be crucial for progression meiosis and differentiation in the planarian germline.

The outcome of m^6^A deposition on RNA in planarians remains to be elucidated and is likely to require transcript-specific analyses. One conserved feature that has been observed in transcriptomic analyses of *S. mediterranea* is the abundance of m^6^A near the stop codon of mRNAs [60, 61]. The presence of m^6^A in the coding region of mRNAs proximal to the stop codon triggers mRNA degradation via a translation-dependent pathway in human cells [76, 77]. Interestingly, Cui et al. [60] detected increased neoblast abundance, decreased levels of m^6^A modification, and increased abundance of *cyclin-dependent kinase 7* (*cdk7*), *cdk9*, and other mRNAs upon *WTAP* knockdown, supporting the idea that destabilization of mRNAs through this post-transcriptional modification is required in the process of cellular differentiation in planarians.

While components of the writer complex and readers are conserved between planarians, *Drosophila*, vertebrates and beyond, erasers seem to have been lost in *S. mediterranea*. AlkB family demethylases, namely FTO and ALKBH5, are conserved between organisms as distant as humans and plants, where they remove m^6^A modifications from RNA. Homologs of m^6^A demethylases were not found in *S. mediterranea* in the current or a previous study [59], suggesting that addition of m^6^A on mRNA is not a reversible process in planarians. Interestingly, ALKBH5 is required for normal testis size and fertility in mice [6], indicating the importance of making m^6^A methylation reversible during spermatogenesis. One possibility is that the loss of dynamic m^6^A regulation was superseded by the continuous cellular turnover that characterizes the homeostatic plasticity of planarian flatworms. Planarians develop, grow, degrow, regenerate, and maintain their tissues by balancing rates of neoblast proliferation and cellular apoptosis [78, 79]. Aside from the possible exception of germline stem cells, which have been reported to give rise to somatic tissues under a specific regeneration experiment [80], the neoblast commitment to differentiate not known to be reversible (see [81]) . Once a neoblast commits to a specific somatic lineage it may no longer have the ability to proliferate, de-differentiate, or slow down the process of differentiation. The same may be true of germline stem cells once these ignite the process of spermatogenesis or oogenesis. In this sense, addition and removal of m^6^A would no longer be an advantageous mechanism to balance proliferation and differentiation of cells, as these cellular processes are regulated at the level of neoblasts proliferation and commitment to differentiation.

The outcome of m6A deposition depends on the molecular function of reader proteins that identify this modification in mRNA. Dagan et al. [59] identified six reader homologs in *S. mediterranea* (one YTHDC homolog and five YTHDF homologs). Of these, only knockdown of the YTHDC homolog resulted in a phenotype in asexual planarians. It is unlikely that the lack of observed RNAi phenotypes for YTHDF homologs is explained by functional redundancy, as these are expressed in limited non-overlapping cell types [59]. Interestingly, our experiments in sexual planarians revealed roles for three different YTHDF readers in germline development. *Smed-YTHDF1-2* expression is enriched in ovaries and crucial for oocyte development (Figure 5E). The mammalian homolog YTHDF1 stimulates translation directly and indirectly via stimulation of eIF3C translation, and amplification of *YTHDF1* stimulates proliferation in ovarian cancer [82]. We hypothesize that Smed-YTHDF1-2 regulates proliferation and differentiation of oogonia by ensuring timely translation of its targets. On the other hand, *Smed-YTHDF2-3* and *Smed-YTHDF2-2* are preferentially expressed in testes and support spermatogenesis (Figure 6, E-H). YTHDF2 is known to recruit the CNOT1/CCR4 complexes to promote deadenylation and decay of its targets [83]. We hypothesize that YTHDF homologs in *S. mediterranea* promote oogenesis progression by degradation of mRNAs that promote stemness at cost of differentiation. Further studies of these specific readers and identification of their targets in *S. mediterranea* will provide insight into the mechanisms by which mRNA regulation via m^6^A modification regulates germline development.

## MATERIALS AND METHODS

### Planarian cultures

A laboratory line of *S. mediterranea* sexual hermaphrodites [84] was used for the bulk of experiments reported in this study with the exception of analyses specifically indicated as performed using asexual animals, in which case a CIW4 clonal laboratory strain [85] was used. Sexual animals were maintained at 18°C in 0.75X Montjuïc salts [86]. Asexual animals were housed in incubators at 21°C filled with 1x Montjuïc salts. Planarians were maintained in plastic containers of ∼2 liter capacity and in the dark except during twice-per-week feedings of beef calf liver, which occurred on benchtops at room temperature. Animal husbandry containers were cleaned after each feeding and replenished with fresh media. Animals were starved for at least one week prior to experimentation or fixation.

### Ablation of stem cells by x-ray irradiation

Asexual planarians were exposed to x-ray irradiation as described in Tasaki et al. [87] minus coverage by lead shields. Total RNA was extracted from irradiated planarians 7 days following irradiation using TRIzol as per the manufacturer protocol (Thermo Fisher Scientific, Waltham, MA) and used for reverse transcription quantitative PCR (RT-qPCR) as previously described [46, 88]. The primers used in RT-qPCR analysis were: CCATGCTTGCAAACAGAAGG and TCCTTGCTCCATTGCTCTTC for *Smed-METTL3*; CTGGTGTGGTAGTGGTGAAG and TGAAATAGTGCACCAGGCTC for *Smed-METTL14*; and CAAGGACAAATGTTGCCTGTA and CAACCCATCGATCCAACTCT for *nanos*. *Smedwi-1* and *ß-tubulin* primers were described in [46, 88].

### Cloning of m6A methyltransferase machinery and YTHDF homolog cDNAs

Partial cDNA sequence clones were generated from PCR-based amplification of reverse transcription products generated from sexual *S. mediterranea* total RNA primed with oligo-(dT) and random primers as per GoScript cDNA synthesis kit manufacturer protocols (Promega, Madison, Wisconsin). PCR using gene-specific primers was performed using Long PCR Master Mix (Promega, Madison, WI) and product were ligated into pGEM-T vectors as per manufacturer instructions for T/A cloning (Promega, Madison, WI). The oligos used for amplification of gene specific sequences are the following:

ATTAATGGGCGATACTTGGAAAG and TCTTTCAACAACTTCTGGATCTACC for *METTL3*; TGGCTGATAACTCAATCAATGATAC and ATTATCGACGAGATCCTCCAGTAG for *METTL14*; GCCCAAATTGAAGACATGAAG and TCCTCATCATCCTCCTCCTC for *WTAP*; CAAATCCATTTTGTGAATCAGG and TCCAGAACATTGGATCGATTAC for *ythdf2-3*; TGGAGTCTTTACCACATTCCTG and AATAATAAAGGCCTCCGCTTC for *ythdf2-2*; TATGCCCCAATTCAGAATCC and CCGCCATAAATGTATCCAATG for *ythdf1-1*; GCTCCATCATTTACGCCAAG and AGGAGGGATACCCCTGTATG for *ythdf2-1*; AAGATCAGCCCAATGAATCG and CAATCACGCGAAAGATATGC for *ythdf1-2*.

### Whole-mount *in situ* hybridization

Whole-mount colorimetric *in situ* hybridization was performed as described by Pearson et al. [89] while FISH and DAPI staining were performed following the protocol described by King and Newmark [90] and Wang et al. [57], respectively. Large animals (∼1.0 cm or larger) were used in *in situ* hybridization analyses of sexual animals, whereas animals of 0.5-0.7 cm were used in analyses of asexual specimens. Digoxigenin-11-UTP (Cat. No 11209256910, Roche Diagnostics GmbH, Mannheim, Germany) and Fluorescein-12-UTP (Cat. No. 11427857910, Roche Diagnostics GmbH, Mannheim, Germany) were used in riboprobe synthesis by *in vitro* transcription using cDNA clone sequences as templates. Low magnification images of colorimetric and FISH samples were obtained using a Zeiss V.16 SteREO microscope coupled to a Canon EOS Rebel T3 camera. For high magnification imaging of FISH and DAPI staining, single confocal plane images were generated from mid-point depth of gonads using a Nikon C2+ confocal microscope with NIS-Elements software.

### Disruption of gene expression by RNAi

Double-stranded RNA was generated by *in vitro* transcription using T7 RNA Polymerase from templates generated by PCR from cDNA clones as previously described [68, 91]. Planarians were fed twice per week with dsRNA 100-200 ng/μL (final concentration) delivered using a 2:1 beef liver puree to water mixture, which was fed to groups of planarians to satiation. Animals were amputated of fixed for analyses by DAPI, immunostaining, and/or *in situ* hybridization a week following the final of five dsRNA feedings (regeneration experiment in asexual planarians) or 8-9 dsRNA feedings (sexual planarians and germline stem cell cluster analysis in asexual planarians).

### Analysis of testis distribution and sperm development

A week following completion of RNAi treatments, samples were fixed and bleached as described above for *in situ* hybridization (minus methanol dehydration and rehydration steps). After bleaching, the samples were washed in PBSTx twice and then incubated in DAPI (ACROS Organics, Morris, NJ) solution (1:1000 dilution of 1mg/ml DAPI stock solution in PBSTx) overnight while rocking at 4°C. After incubating overnight, the samples were washed 4 times with PBSTx and then mounted on slides with 4:1 glycerol:PBS and imaged under UV light with a Zeiss V.16 SteREO microscope equipped with a Canon EOS Rebel T3 camera (for low magnification) or a Nikon C2+ confocal microscope using a 20X objective and running NIS Elements C software (for high magnification).

### Analysis of asexual planarian regeneration by immunofluorescence and DAPI staining

Asexual planarian subjected to RNAi treatments were amputated anterior and posterior to the pharynx a week following the last dsRNA feeding. Then samples were fixed and analyzed using DAPI and anti-SYNORF1 (1:250 dilution; clone ID: 3C11, Developmental Studies Hybridoma Bank, Iowa City, IA) as described previously [92].

## ACKNOWLEDGEMENTS

This work was supported by a Japanese Society for the Promotion of Science Overseas Research Fellowship to JT and by the National Institutes of Health [Award No. R15 HD082754] to LR. The authors declare no conflicts of interest.

