## Supplementary Material Legends for "Contributions of m6A RNA methylation to germline development in the planarian *Schmidtea mediterranea*"

**SUPPLEMENTARY FIGURE LEGENDS**

**Supplementary Figure S1. Neoblasts and GSCs were not affected by perturbation of $m^6A$ methylation pathway component expression in asexual planarians.** Neoblasts (marked by *gH4*) and presumptive germline stem cells (marked by *nanos and gH4*) were present at comparable levels in control RNAi samples and after more than four weeks of $m^6A$ writer and reader gene knockdown. Fraction of samples showing phenotypes undistinguishable from controls is shown in parenthesis. Scale bar: 50 μm.

**Supplementary Figure S2. No defects in planarian regeneration were observed upon $m^6A$ methyltransferase gene knockdown. (A-B)** Normal regeneration is observed in trunk (A) and tail (B) fragments of planarians subjected to RNAi-mediated disruption of *Smed-METTL3* expression, *Smed-METTL14* expression, or *Smed-METTL3;Smed-METTL14* simultaneous knockdown. DAPI staining of cell nuclei (gray) is shown along with visualization of the brain and ventral nerve cords labeled using anti-SYNORF antibodies (green). Brackets show regenerated head or tail. Asterisk indicates position of the pharynx. Scale bar is 500 μm.

**Supplementary Figure S3. No decrease in fission events observed during $m^6A$ methyltransferase gene knockdown.** Graph portraying cumulative fission events observed during knockdown of writer complex genes (*METTL3;METTL14(RNAi)* or *WTAP(RNAi)*) and reader genes (*YTHDF1-1(RNAi)*, *YTHDF2-3(RNAi)*, *YTHDF2-2(RNAi)*, *YTHDF1-2(RNAi)*, or *YTHDF2-1(RNAi)*).
