## Supplementary figures and images for "Contributions of m6A RNA methylation to germline development in the planarian *Schmidtea mediterranea*"

### Supplementary Figures

Tasaki_Fig_S1

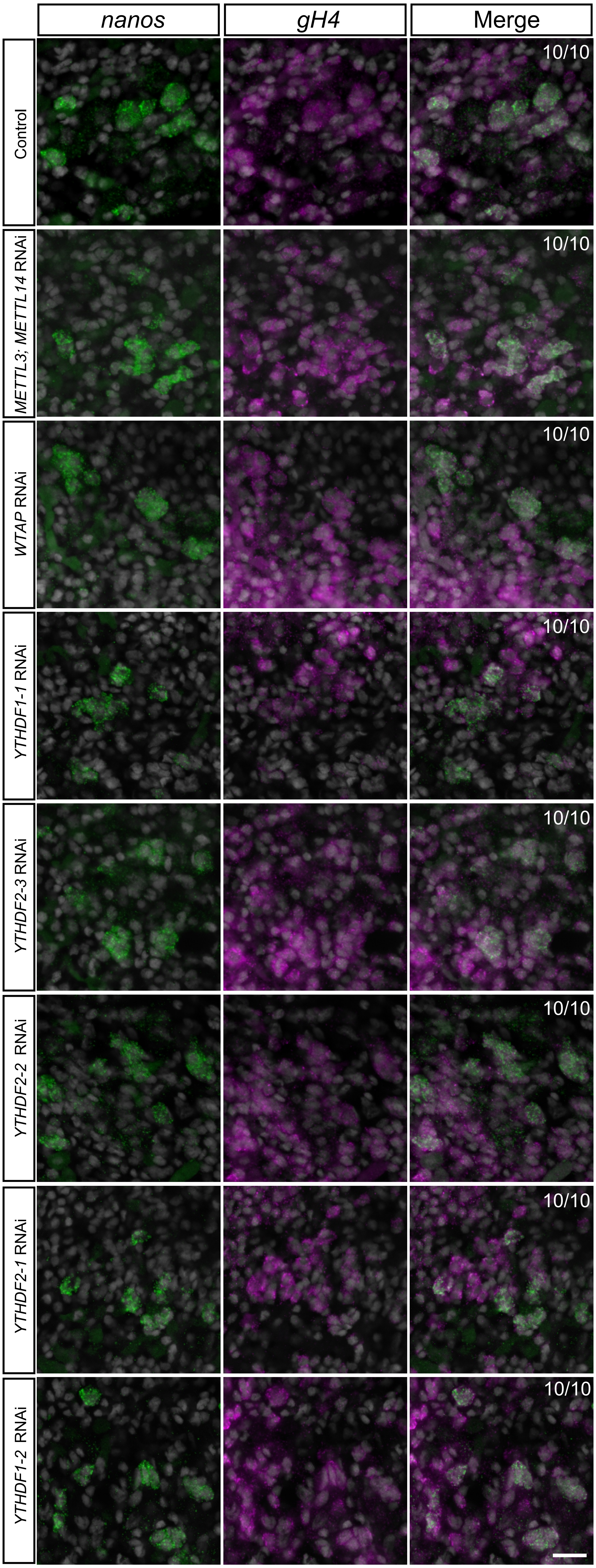

Tasaki_Fig_S2

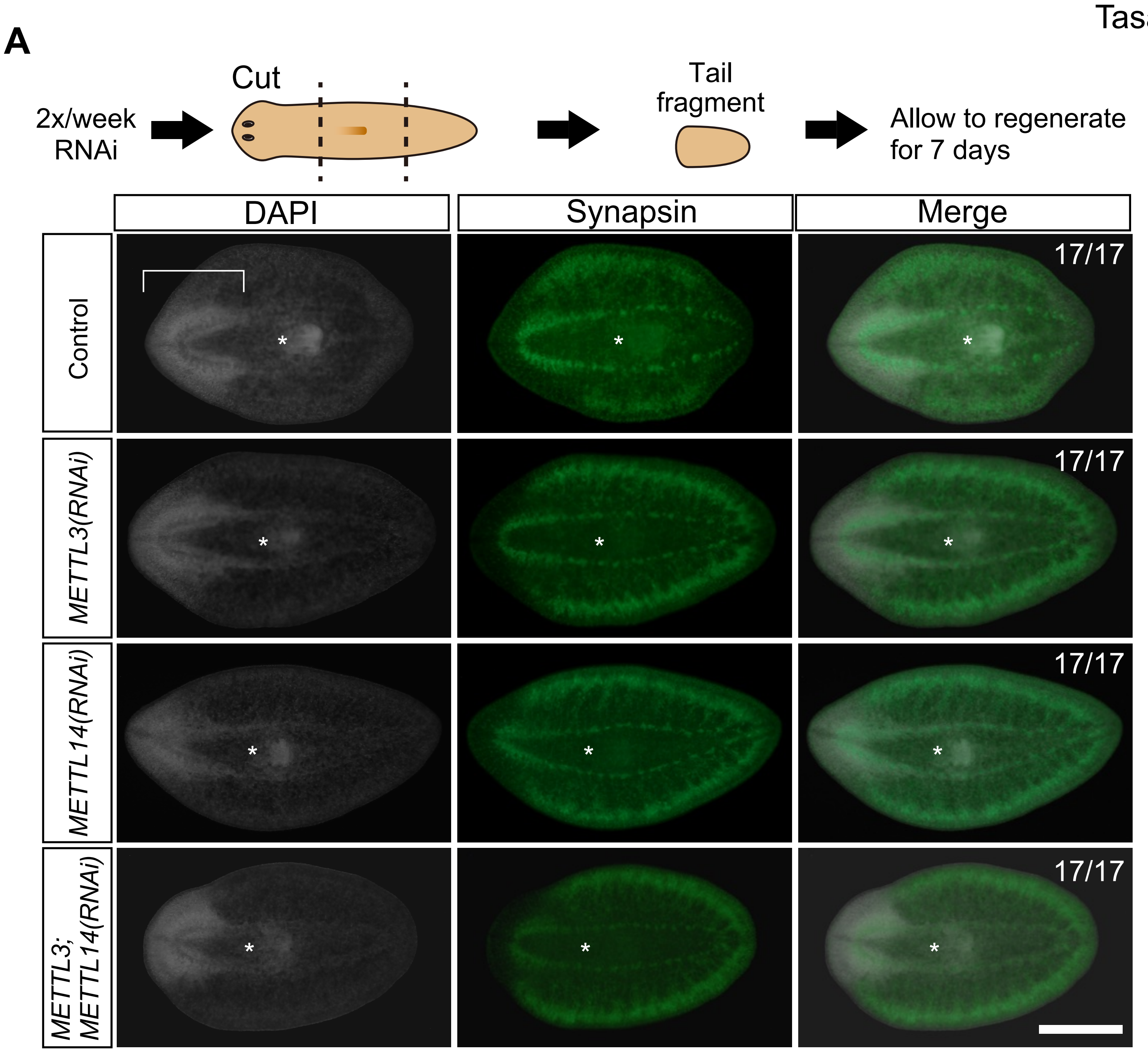


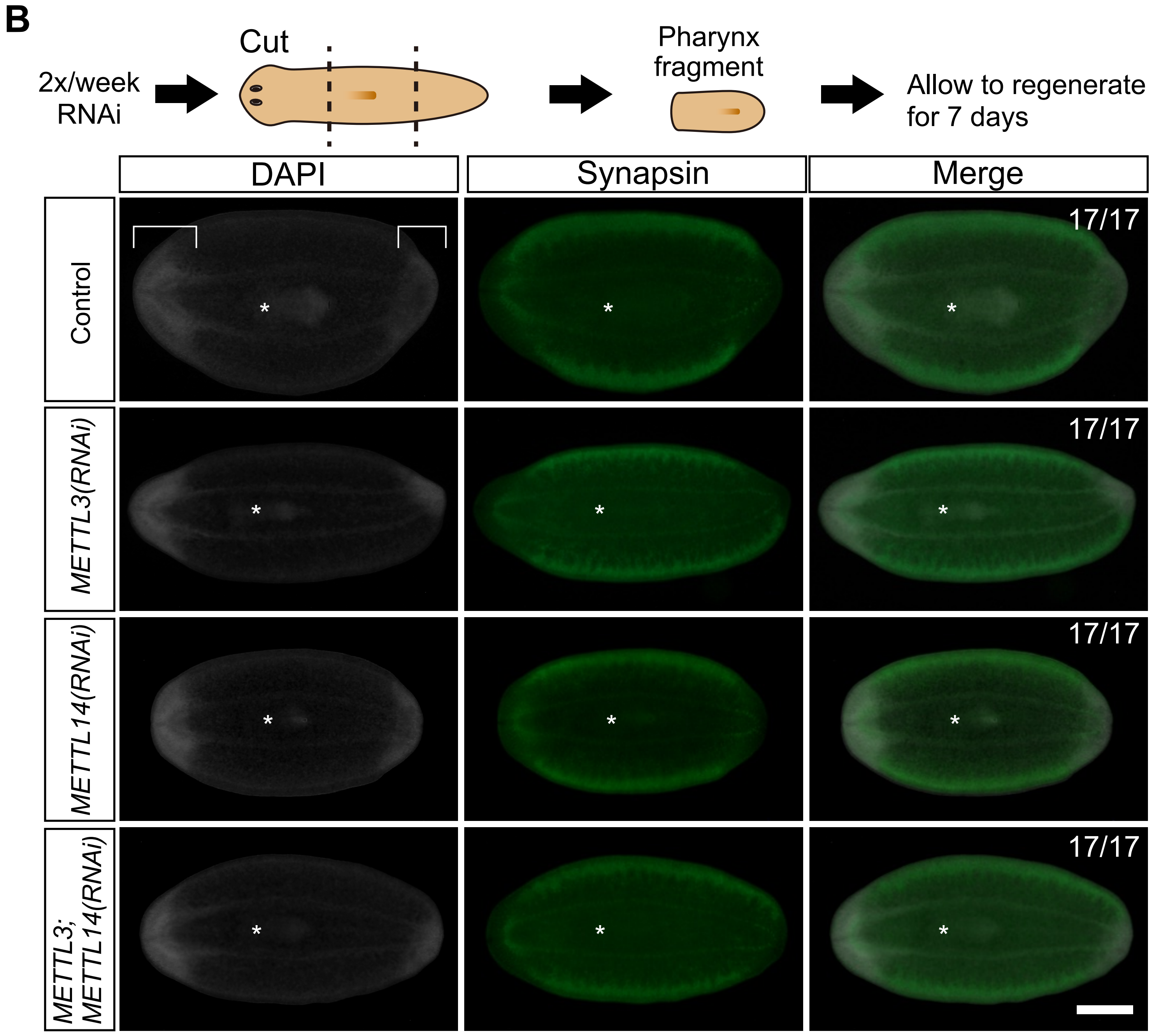

Tasaki_Fig_S3

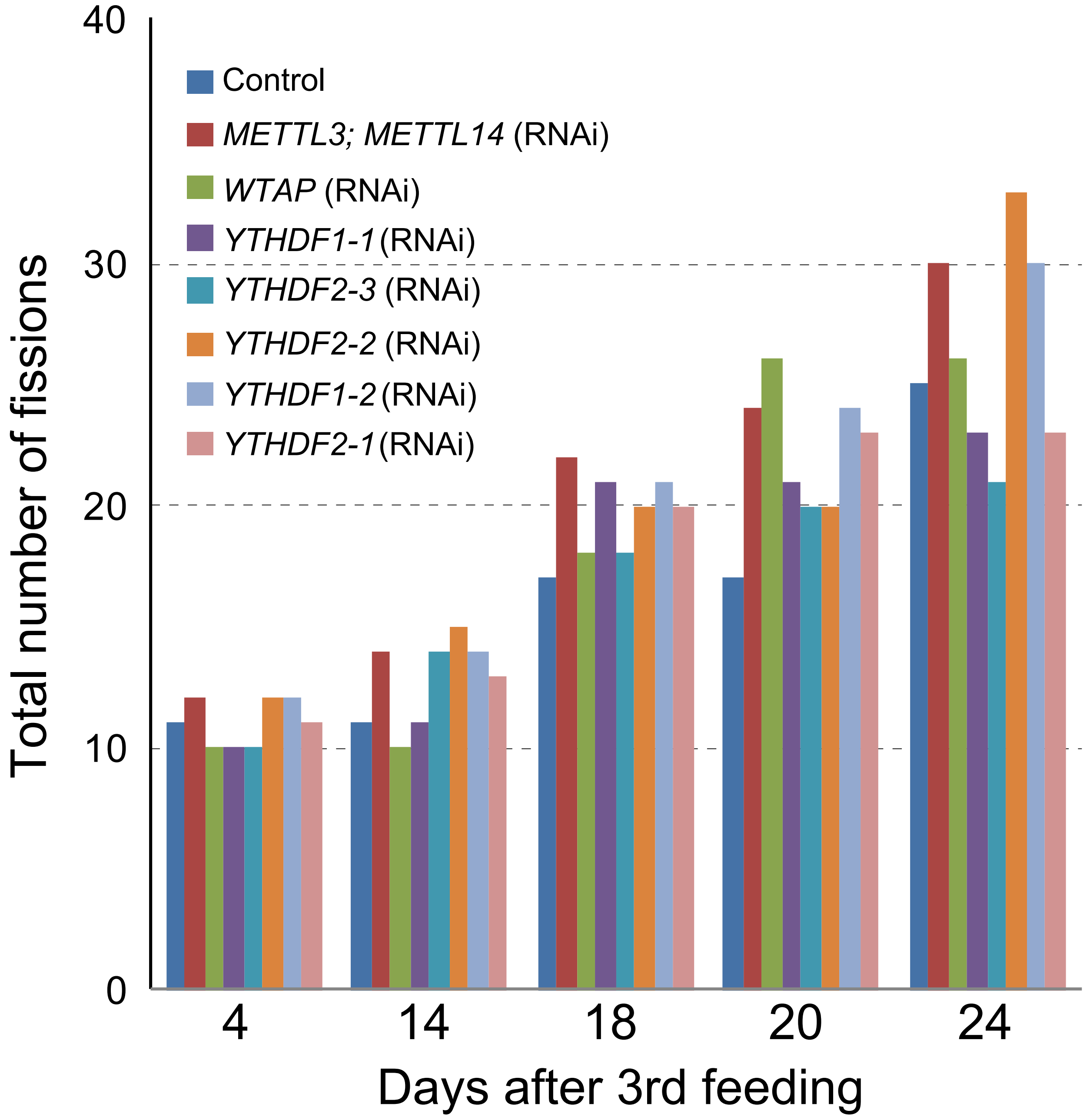
